# Microscopy-informed structural connectivity mapping in the in vivo human brain via domain adaptation

**DOI:** 10.64898/2026.06.14.732211

**Authors:** Silei Zhu, Nicola K. Dinsdale, Saad Jbabdi, Karla L. Miller, Amy F.D. Howard

## Abstract

Characterising human brain connectivity remains a major challenge in neuroscience. Multimodal datasets combining diffusion MRI with high-resolution microscopy in the same brain offer a unique link between macroscopic imaging and microstructural detail, but we lack tools to leverage these data to improve connectivity estimates for in vivo human imaging.

We present a deep learning model that predicts high-resolution microscopy-informed fibre orientations from diffusion MRI. The model uses microscopy-derived three-dimensional fibre orientation maps as biologically grounded training targets. It is trained on a bespoke macaque dataset integrating in vivo MRI, postmortem MRI, and whole-brain microscopy, and then translated to in vivo human imaging. We use domain adaptation to predict fibre orientations from diverse MRI datasets: first to bridge differences in tissue state in the macaque (postmortem to in vivo), and then to generalise across species (macaque to human).

Our method derives microscale-informed fibre architecture from diffusion MRI without requiring microscopy at inference. It leverages data that can easily be acquired only in animal models whilst generalising to in vivo human diffusion MRI with minimal acquisition requirements. The microscopy-informed fibre orientation distributions support biologically meaningful tractography, enhancing superficial white matter and cortical-subcortical pathway delineation for in vivo human data. More broadly, this work establishes a general framework for transferring microstructural information from microscopy to non-invasive imaging, enabling biologically informed mapping of brain connectivity.

## Introduction

Acquiring datasets that link MRI to microscopy in the same brain has enabled more precise methods for probing the mesoscopic organisation of brain networks. While diffusion MRI (dMRI) offers a non-invasive method to estimate brain connectivity^1,2^, its accuracy is limited by spatial resolution and reliance on signal modelling, especially in regions with complex fibre architecture^3–6^. In contrast, microscopy provides detailed maps of fibre pathways, but is limited to postmortem tissue and is typically acquired in non-human species (e.g. rodents and non-human primates)^7–9^. As multimodal datasets linking MRI and microscopy become increasingly available, new opportunities are emerging to integrate these complementary perspectives^8,10^. A key methodological challenge is how best to harness such dMRI-microscopy ex vivo data to inform and enhance in vivo dMRI brain mapping. Here, we present a deep learning framework trained on paired dMRI and myelin-sensitive microscopy data in the macaque brain, which learns to predict fibre orientations from dMRI-only datasets.

Deep learning has shown promise for improving fibre orientation estimation from dMRI^11,12^. One of the primary challenges with dMRI is the requirement for computational models to infer fibre orientations from MR signals. These models typically require strong assumptions, such as representing axons as sticks^13^ or assuming the same diffusion signal (“response function”) for fibres across the white matter^14^. Inaccurate or biased fibre orientation reconstruction can lead to errors in downstream tractography^13–15^.

Recently, deep learning approaches have offered a data-driven alternative that eliminates the need for explicit biophysical modelling^12,16^. However, a key challenge in deep learning methods is generalisability to datasets that do not match the distribution of the training data^17^. This constraint poses difficulties when translating models across imaging modalities (microscopy to MRI), tissue states (postmortem to in vivo), and species (rodent or monkey to human), especially given that most existing dMRI-microscopy datasets are derived from postmortem non-human brains^8,18–21^. Substantial differences exist between dMRI data acquired from postmortem and in vivo tissue: while postmortem dMRI allows for higher spatial resolution and superior data quality due to longer scan times, fixed tissue exhibits reduced diffusivity and altered relaxation times^22^. Moreover, models trained on species with simpler brain architecture, such as rodents, may not generalise well to humans that have highly folded brains and complex white matter architecture (e.g., multiple fibre populations in a single voxel). These disparities suggest a data distribution mismatch that may cause models trained exclusively on postmortem non-human dMRI to produce unreliable results when applied directly to in vivo human data.

Domain adaptation in deep learning provides a potential solution by enabling neural networks to become invariant to differences in data distributions across domains, thereby mitigating the effects of dataset heterogeneity (Figure 1b, “Domain adaptation”). Domain adaptation has previously been used in MRI to design networks that generalise across different imaging centres, scanners, and acquisition protocols^23–25^. Here, we apply this concept to develop a network that predicts microscopy-derived fibre orientations from dMRI, while remaining invariant to tissue and species differences between in vivo/postmortem macaque and human data^8^. This approach enables training with paired MRI-microscopy data from a macaque brain, allowing the network to benefit from high-quality postmortem data, while ensuring generalisability to in vivo human applications (Figure 1a).

**Figure 1:**
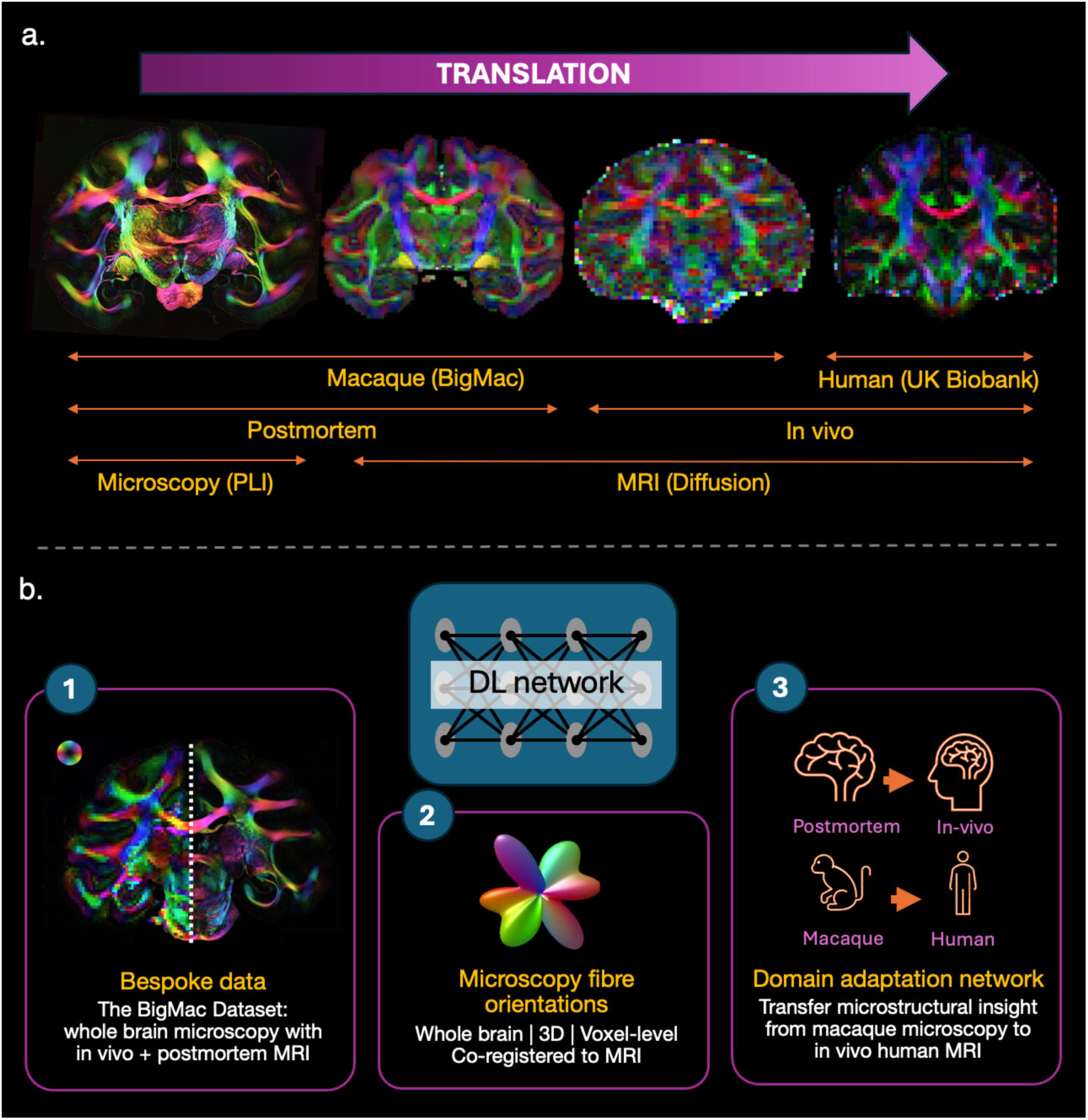
Translating microscopy insights to in vivo brain connectivity imaging. a) Our method enables the transfer of microstructural information from postmortem macaque brains to in vivo human neuroimaging. This is achieved by bridging three critical domains: (i) imaging modality, moving from microscopy (polarised light imaging (PLI)) to diffusion MRI (dMRI); (ii) tissue state, translating from postmortem to in vivo; and (iii) species, adapting features learned from the macaque brain (BigMac dataset) to the human brain (e.g., UK Biobank). b) To enable transfer of microscopy-derived microstructural information to in vivo human imaging, our method involves three key components: (i) bespoke training data consisting of whole-brain microscopy paired with in vivo and postmortem MRI; (ii) 3D microscopy-informed fibre orientations to act as a “ground truth” during training; and (iii) a domain adaptation network to translate insights across diverse datasets and enable transfer of microscopy insights from postmortem macaque data into the living human brain.

For optimal performance, our network requires training data that satisfy three criteria (Figure 1b, “Bespoke data”). First, the microscopy and dMRI data must be acquired from the same brain, with microscopy providing “ground truth” fibre orientations during training. Using data from the same subject avoids microstructural mismatches due to inter-subject variability, which would otherwise hinder accurate MRI-microscopy mapping. The microscopy and dMRI data must nonetheless be registered to each other with high accuracy, which presents a challenge even when they come from the same brain. Second, the dataset should include in vivo and postmortem MRI from the same brain (again to avoid confounding inter-subject variability). Postmortem dMRI serves as an intermediary between in vivo dMRI and postmortem microscopy, typically offering higher spatial resolution and signal-to-noise ratio (SNR) than in vivo MRI, which may better facilitate learning of the MRI-microscopy relationship. Third, to limit the domain gap with human data, the training set should be derived from tissue with structural complexity comparable to the human brain, favouring higher primates over rodents. Crossing fibres are reported to be present in over 80% of white matter voxels in humans; in contrast, rodent brains contain relatively little white matter and exhibit simpler fibre architecture, with far fewer multi-fibre voxels^26^. Networks trained on rodent data may therefore struggle to generalise well to human applications. The macaque brain, by contrast, shares many key features with humans, including substantial cortical folding and complex white matter organisation.

We trained our network using the BigMac dataset^8^, an open data resource that provides co-registered in vivo dMRI, postmortem dMRI and high-resolution microscopy (including polarised light imaging, PLI) from the same macaque brain, thereby satisfying the criteria outlined above (Figure 1b). The trained network takes either in vivo or postmortem dMRI as input and outputs a predicted microscopy-informed fibre orientation distribution (FOD) for each voxel, characterising the local three-dimensional fibre orientation structure.

During training, microscopy-derived fibre orientation distributions are used as ground truth (Figure 1b, “Microscopy fibre orientations”). These are derived from BigMac polarised light images, which provide fibre orientations at a resolution of 4 μm per pixel from thin tissue sections with 50 μm thickness sampled every 350 μm across the brain. We derive “hybrid” 3D orientations that combine the high-resolution in-plane 2D orientations from PLI with out-of-plane orientations from dMRI^27^. Specifically, the dMRI data are processed to estimate a distribution of orientations in each large dMRI voxel (the “ball-and-sticks model”)^13^. We then identify the 3D “stick” from this distribution with the closest in-plane match to the 2D PLI orientation. Its inclination angle is then combined with the in-plane orientation from PLI to provide a final hybrid 3D orientation at the PLI spatial resolution (Figure 1b). These 3D orientations are then aggregated across the local neighbourhood of a voxel to generate a “hybrid MRI–microscopy FOD” (one hybrid FOD per dMRI voxel), which serve as the training target.

As PLI is a myelin-sensitive signal, training against the hybrid PLI-MRI orientations encourages the model to learn more myelin-specific information^27^. Our previous work has shown how these hybrid orientations differ from those obtained using standard dMRI, as the inclusion of microscopy information drives them to exhibit distinct qualities such as lower dispersion and fewer noisy crossing fibres, particularly near and within the cortex, which in turn improves fibre tracking accuracy. The aim in the present work is to translate these “microscopy-derived benefits” to in vivo applications by training the network on hybrid outputs to retain these advantageous properties in living participants.

During inference, the trained network generates microscopy-informed FODs from new dMRI data without requiring concurrent microscopy. We first validate our trained network using held-out in vivo macaque data and then retrain and extend the model to in vivo human MRI. We demonstrate the extent to which microstructural information can be translated across tissue states and species, from postmortem macaque to living humans, allowing microscopy-derived “ground truth” to inform the fibre orientation predictions from MRI alone. By bridging microscopy and MRI, our framework aims to transfer microstructural insights from microscopy to in vivo imaging for more accurate brain connectivity mapping in both research and clinical settings.

## Results

### Design of the network architecture

We developed a domain adaptation network with three key components: a feature extractor, a predictor, and a domain discriminator (Figure 2). The network was designed to predict microscopy-informed FODs from dMRI whilst mitigating the domain shifts across datasets (postmortem versus in vivo dMRI, or macaque versus human). Specifically, the feature extractor and predictor generated FODs from input dMRI, while the domain discriminator enforced invariance across dMRI domains with different distributions. The network took a 3×3×3 cube of dMRI voxels as input (rather than individual voxels) to include local neighbourhood information, which was combined through the convolutional layers. For each cube of dMRI input, the network output a single FOD for the central voxel. During training, this was compared to the hybrid dMRI-microscopy FODs, which provided a microscopy-informed “ground truth”. Both the dMRI input and FOD output were represented using spherical harmonics. Encoding the dMRI signal in this form allows the network to generalise across datasets without strict acquisition requirements on the number or orientation of diffusion-encoding gradients: the number of gradients only needs to meet or exceed the number of spherical harmonic coefficients used. The network requires only single-shell dMRI (data acquired at a single b-value), facilitating wide application across existing and future datasets. The spherical harmonic FOD output is a common format and compatible with existing tractography algorithms for downstream connectivity mapping. We call the method “MiFO” for Microscopy-Informed Fibre Orientations.

**Figure 2.**
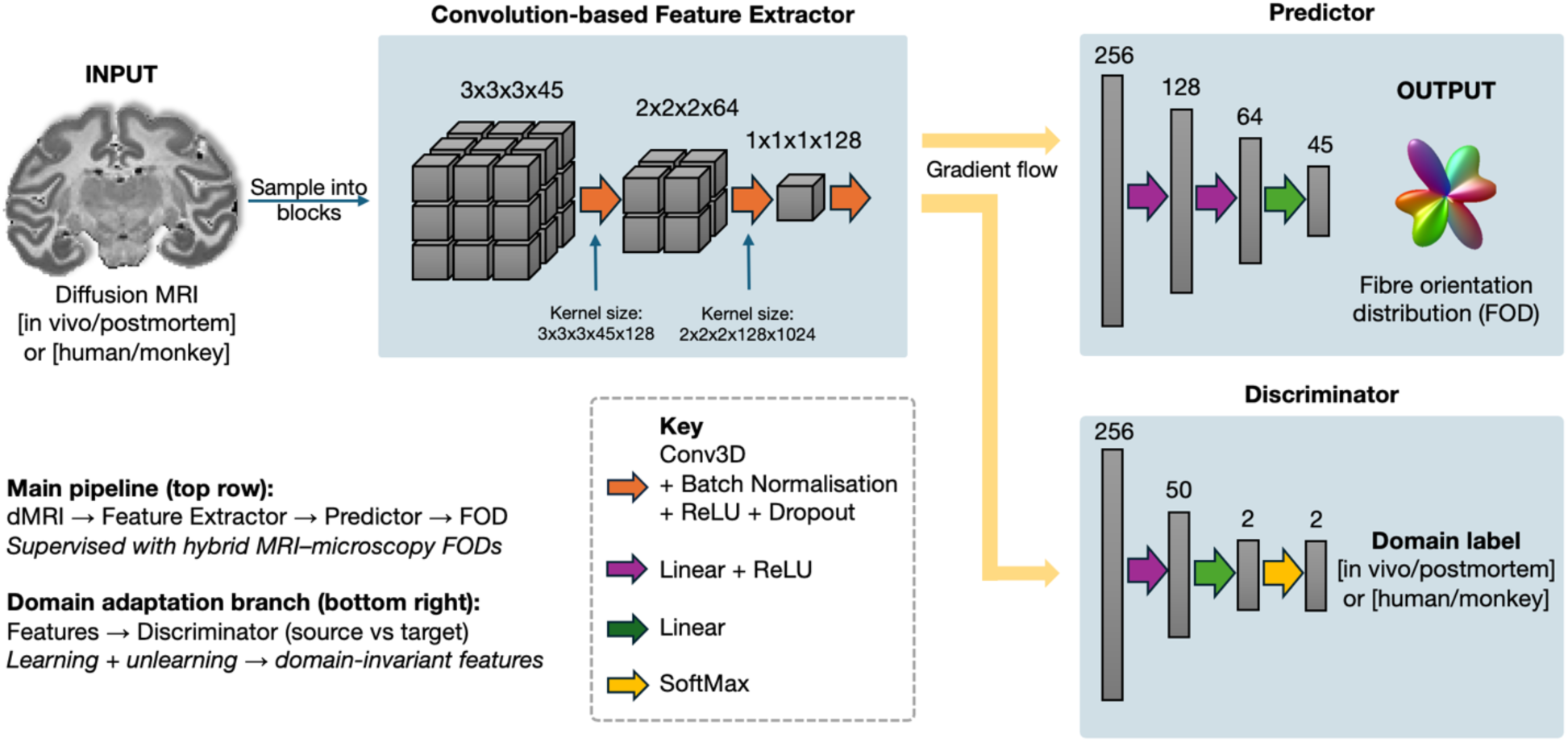
Network architecture. The proposed domain adaptation network estimates microscopy-informed fibre orientations (MiFO) from dMRI signals acquired both postmortem and in vivo. The input dMRI is first sampled into blocks and passed through the network. The architecture consists of three main components: a feature extractor, a predictor, and a domain discriminator. The feature extractor and predictor map dMRI inputs to FOD outputs and are trained using hybrid MRI–microscopy FODs as ground truth supervision. Both the input dMRI and output FOD are represented using 45 spherical harmonics. The domain discriminator encourages domain invariance by distinguishing between i) postmortem and in vivo MRI inputs during initial training on macaque data, and ii) macaque and human MRI during human adaptation. The discriminator outputs a softmax probability over domain labels (in vivo/postmortem or human/monkey). Training consists of two stages: learning and unlearning. In the learning stage, the feature extractor and predictor are trained to estimate FODs, while the discriminator learns to distinguish between domains. In the unlearning stage, the discriminator is fixed while the feature extractor is optimised to confuse it. This encourages the learning of domain-invariant representations.

The network was trained following a two-stage adversarial domain adaptation approach inspired by Dinsdale et al., 2021^25^. With in vivo and postmortem MRI from the BigMac dataset as input, the first stage jointly optimised the feature extractor and predictor for the primary task of microscopy-informed FOD estimation, while a discriminator was trained to distinguish latent features across input domains (postmortem versus in vivo dMRI). In the second stage, the discriminator parameters were frozen, and the feature extractor and predictor were adversarially optimised to maximise domain confusion whilst maintaining accurate FOD estimation. This process forced the model to learn domain-agnostic features – i.e., features that generalise across postmortem and in vivo data – that represent the shared underlying microstructure.

We then generalised the model to human data using an unsupervised domain adaptation framework, in which labelled macaque data (with MRI and microscopy) provided supervision and unlabelled human data (MRI only) guided the learning of species-invariant features. Following training, the network was applied to in vivo human data to generate microscopy-informed FODs without requiring matched microscopy data.

### Network-reconstructed fibre orientations follow neuroanatomy in both macaque and human

We first applied the network to postmortem (Figure 3a) and in vivo macaque data (Figure 3b) that were not used during training. The postmortem test data were from the same subject as the training dataset but were acquired in a separate session, testing generalisation across acquisitions. The in vivo test data were from a different macaque, testing generalisation across subjects. The same network architecture was subsequently adapted for in vivo human data using unsupervised domain adaptation, with FOD estimation supervised on the macaque data only, while the human data were aligned to the macaque domain via the adversarial training. Here, we evaluated the method on two large open-source dMRI datasets: the Human Connectome Project (HCP)^28^ (Figure 3c) and the UK Biobank^29^ (Figure 3d). These datasets provide complementary benchmarks, spanning high-resolution, high-quality acquisitions with long scan times (HCP: 1.25 mm isotropic resolution, multi-shell acquisition, ∼1 hour scan time in young adults) and shorter-duration, good-quality clinical-style acquisitions (UK Biobank: 2 mm isotropic resolution, multi-shell acquisition, ∼7 minutes in older adults). The HCP represents a data quality ceiling, whilst the UK Biobank demonstrates the network’s utility in more “typical” protocols with shorter acquisition times.

**Figure 3.**
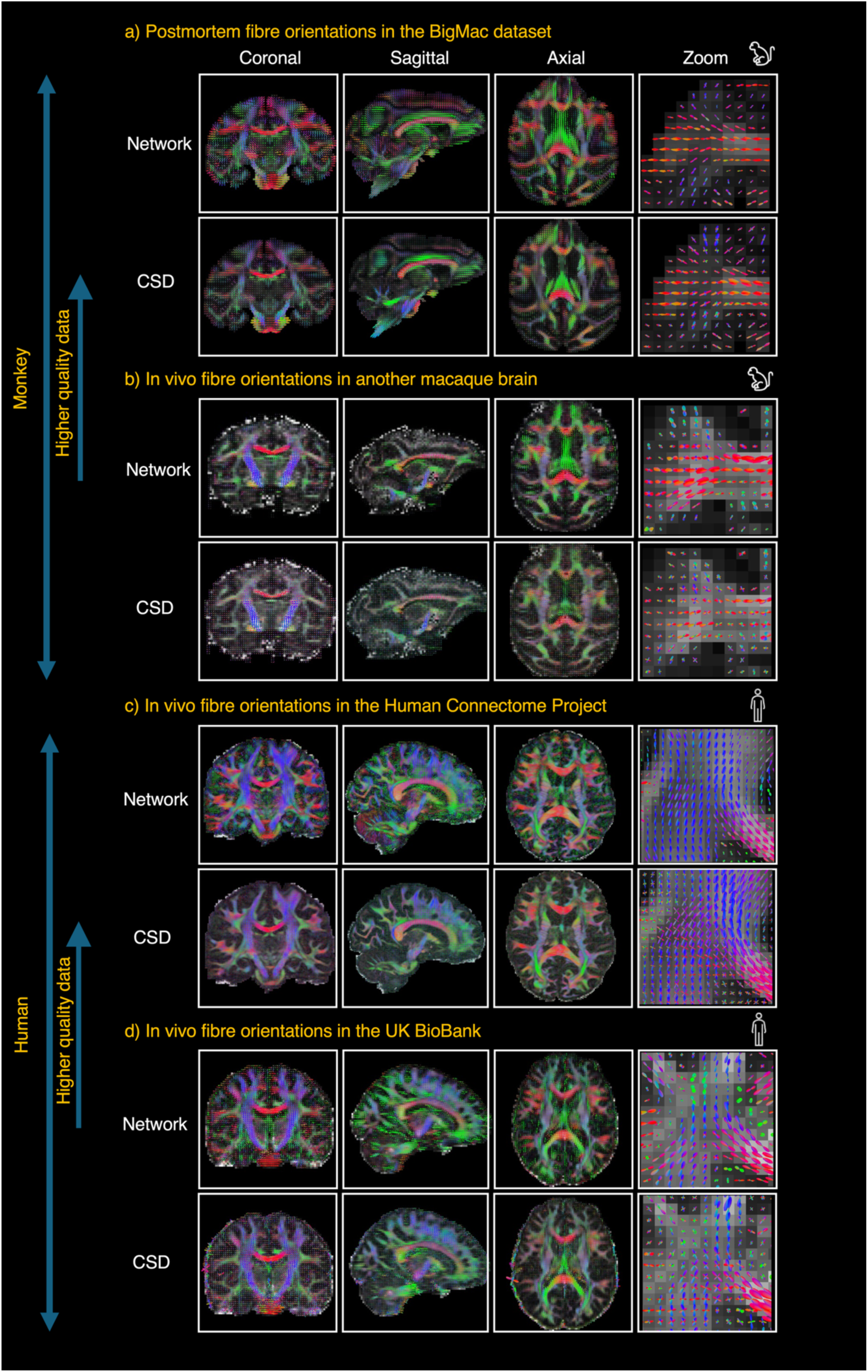
Performance of the microscopy-informed network across tissue types and species. Whole-brain fibre orientation distributions (FODs) estimated by the network and conventional CSD show similar large-scale neuroanatomical organisation. Reconstructions are shown for: a) the postmortem macaque BigMac dataset comprising high-quality data used during training; b) in vivo macaque data acquired on another macaque brain with the same protocol as BigMac, demonstrating cross-subject performance on more challenging, in vivo data; and cross-species translation of the network to in vivo human data, including c) high-resolution HCP data (top research quality, 1hr protocol), demonstrating high-quality human translation and d) lower-resolution UK Biobank data acquired with a more clinical-style protocol (∼7min), demonstrating applicability to typical acquisitions.

In each subject, we produced whole-brain maps of network-predicted FODs, covering the white matter, grey matter, and subcortical regions. The network-predicted FODs in all datasets follow established neuroanatomical expectations, exhibiting primary fibre orientations that are broadly consistent with those derived from conventional diffusion-only constrained spherical deconvolution (CSD) (Figure 3). Both methods recover the expected fibre anatomy, with FODs consistent with all the major white matter pathways such as the easily recognisable corpus callosum, corticospinal tract, and cingulum bundle across all datasets. We also observe noticeable differences between network and CSD FODs, including fewer crossing fibre populations and more “coherent” fibre organisation near the cortex. A systematic comparison, including how network FODs reproduce microscopy features, is presented in Figure 5 below.

We further evaluated whether the network-estimated FODs facilitated anatomically meaningful tractography, a key validation criterion for assessing FOD accuracy. Our method successfully reconstructed major white matter tracts in all datasets. Figure 4a presents example reconstructions of major pathways in humans using the network FODs, including the forceps minor, inferior fronto-occipital fasciculus, and corticospinal tract, demonstrating successful reconstruction of example commissural, association, and projection fibre bundles. The same tracts reconstructed with CSD FODs are provided in supplementary Figure 1. Both approaches reliably reconstructed the tract cores and showed broadly similar trajectories within the deep white matter. Most variation was observed in the cortical streamline terminations, where the network FODs yielded more anatomically coherent results (detailed in Figure 4b below).

**Figure 4.**
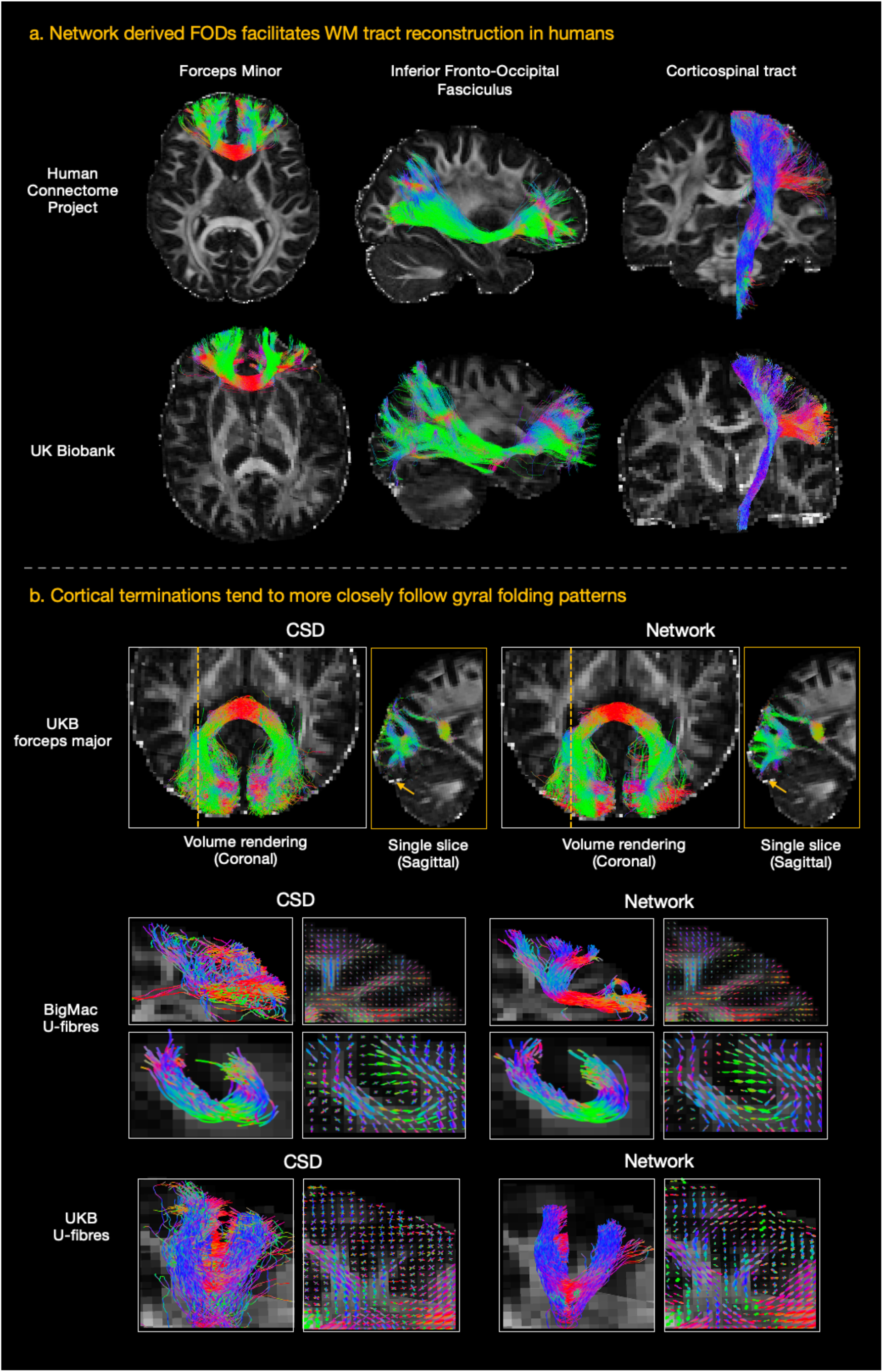
White matter tractography with network and CSD FODs. a) Tractography performed using network FODs successfully reconstructs major pathways in both HCP and UK Biobank human datasets. Example tracts include the forceps minor, inferior fronto-occipital fasciculus, and the corticospinal tract. b) Comparison of cortical terminations with conventional constrained spherical deconvolution (CSD). Reconstruction of the forceps major in the UKB dataset: compared to conventional CSD analysis (left), the network FODs (right) yield more anatomically “precise” cortical terminations (with tracts displayed in 3D coronal view and 2D single slice sagittal view) and improved U-fibre reconstructions in both macaque and human data.

A notable difference was often observed in fibre projections approaching and entering the cortical grey matter. In most cases, streamlines generated from the network FODs appeared to more closely follow cortical folding patterns, producing spatially localised streamlines that remained more confined to superficial white matter rather than tracking spuriously into grey matter. For example, in the forceps major (Figure 4b), the network tended to resolve more anatomically coherent cortical projections that better followed the gyral patterns compared to CSD. Motivated by this observation, we constructed U-fibres, where the network FODs often but not universally more clearly delineate the pattern of fibres curving around the sulcus compared to CSD (Figure 4b). The improved cortical projection patterns are likely due to orientations with high spatial continuity and smoothness at the cortical boundary, as observed in both postmortem macaque and in vivo human data and discussed below (see also Supplementary Figure 2).

### Network FODs reproduce microscopy features in vivo

Although the overall anatomical organisation is consistent between the network and CSD methods, notable FOD differences are also observed (Figure 5, with details in Supplementary Figures 3 and 4). Interestingly, these differences appeared to reflect features of the microscopy data used during training, suggesting successful transfer of microscopy-informed structural information.

**Figure 5.**
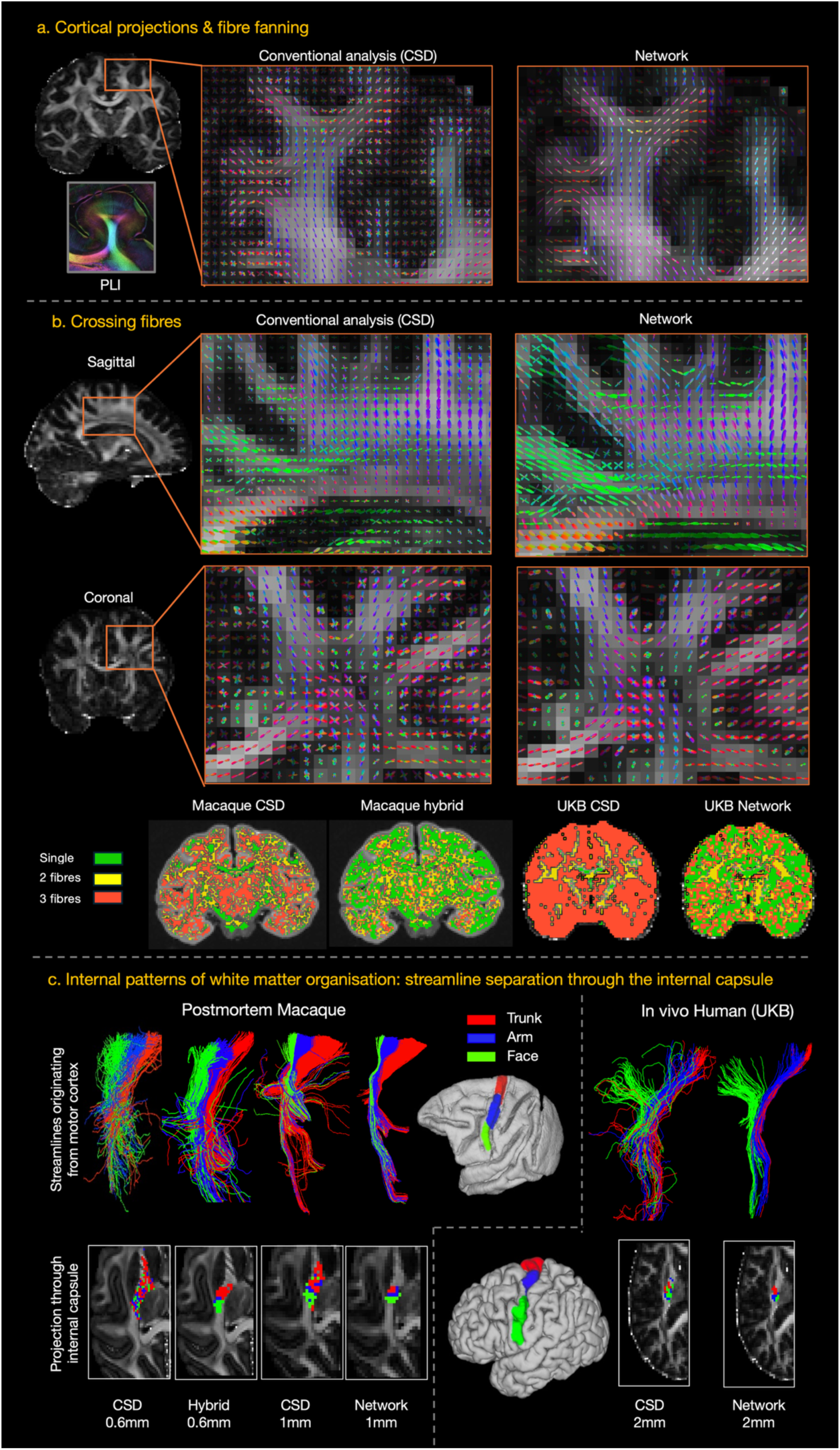
Network FODs reproduce microscopy features in vivo. a) FODs near and into the cortex demonstrate the network’s ability to more clearly resolve fibre fanning patterns observed in myelin-sensitive microscopy, as illustrated in the PLI. b) Although crossing-fibres are present in both methods, the network often resolved fewer crossings than CSD. Crossing-fibre maps were generated by converting FODs to discrete peaks and counting the number of distinct fibre orientations per voxel. These quantitative maps show that the prevalence of crossing fibres in the network outputs mirrors that of BigMac hybrid FODs on which the network was trained on. c) Reconstruction of motor cortex projections originating from the trunk (red), arm (blue), and face (green) and passing through the internal capsule. In both macaque (postmortem) and human (UK Biobank) datasets, network FODs resolve the expected anterior–posterior topographical organisation within the internal capsule, whereas CSD shows greater mixing of streamlines in this anatomical bottleneck. Streamlines are shown in 3D, with “winner takes all” streamline density map displayed below.

Some of the largest FOD differences were observed in fibres projecting into and through the cortical grey matter (Figure 5a). FODs from conventional CSD analysis often appear noisy, with complex crossing configurations that do not correspond well to underlying myeloarchitectural fibre patterns^14^. This arises because the cortical diffusion signal is sensitive not only to axons, but also to cell bodies and dendrites, resulting in reduced diffusion anisotropy compared with white matter. Combined with partial volume effects near the grey-white matter boundary, this can impair reconstruction of the fibre fanning patterns seen in myelin microscopy, where fibres spread from deep white matter into the cortex. A downstream consequence is the well-known “gyral bias” in tractography, whereby streamlines preferentially terminate at gyral crowns rather than sulcal walls or fundi, potentially leading to inaccurate estimates of cortical connectivity. This effect can be partially mitigated by increasing the MRI resolution^20^, with more consistent gyral fanning patterns found in humans in the higher-quality HCP data (Supplementary Figure 4), compared to the lower-resolution UK Biobank (Supplementary Figure 3). In contrast, the network FODs more closely captured coherent fibre trajectories at the grey-white matter boundary, with fibres fanning into the cortex in agreement with microscopy. The network recovered predominantly radial cortical orientations consistent with expected myelinated fibre architecture, with fibre fanning patterns across the gyrus more closely matching both the 2D PLI data and the hybrid orientations used during training. This suggests successful transfer of myelin-specific microstructural information from microscopy to in vivo MRI and indicates that tractography derived from the network FODs may be more specific to myelinated axons.

The network-derived FODs also exhibit lower dispersion (more coherently aligned fibre bundles) and fewer crossing-fibre voxels than conventional CSD (Figure 5b). To determine whether this reflected a network artefact or a learned property of the training data, voxels were quantitatively classified according to whether they contained one, two, or three fibre populations. Comparisons across the original macaque hybrid orientations, the network prediction in humans, and conventional CSD in both species showed that the prevalence of crossing fibres in the network-derived human FODs closely matched that of the BigMac hybrid FODs used during training. This suggests that the network FODs in the human data reproduce the patterns of the hybrid method and capture microscopy-informed structure, rather than artificially suppressing crossing fibres.

Whether these characteristics fully reflect biological ground truth remains uncertain and warrants further validation, as microscopy itself is not an unambiguous reference standard. Previous work has shown that estimated fibre dispersion and crossing prevalence vary across microscopy techniques^27,30^. For example, PLI typically shows fewer crossings and lower dispersion than structure tensor analysis of myelin-stained (Gallyas silver) sections from the same BigMac dataset^27^. Nonetheless, the network successfully learns microstructural features from microscopy (in another species) and transfers them to in vivo human MRI.

Figure 5c demonstrates a further example of translating microscopy-informed features across datasets. The topographical organisation of motor cortex projections within the internal capsule is well established, with trunk, arm and face representations arranged along the anterior-posterior axis and fibres originating from more medial cortical regions projecting more anteriorly^31,32^. In our previous work^27^, microscopy-informed FODs in the BigMac dataset were able to resolve this known somatotopic organisation within the internal capsule, though this structured anterior-posterior organisation was not clearly resolved using conventional CSD.

Here, we show that the network recreates this pattern in both unseen postmortem macaque data and human UK Biobank data. As shown in Figure 5c, conventional CSD tractography fails to clearly differentiate tracts originating from distinct motor cortex regions, leading to mixed streamlines within the internal capsule, a known bottleneck region for tractography. In contrast, tractography derived from the network-predicted FODs demonstrates a clearly separable anterior-posterior organisation, closely resembling the pattern previously observed using the hybrid dMRI-microscopy approach in postmortem data.

To quantitatively evaluate the network’s performance against an anatomical ground truth, we compared network-derived tractography to a weighted connectivity matrix obtained from retrograde tracer data in the macaque (Supplementary Figure 5a)^33–35^. The network outputs demonstrated comparable performance to conventional CSD in predicting tracer-based connectivity, with a similar area under the ROC curve (AUC = 0.916 for the network and 0.911 for CSD), and a reduced false positive rate for the network at a fixed threshold of 0.0001 (Supplementary Figure 5b, Top). Correlations between diffusion MRI- and tracer-derived connectivity were also similar across all reconstruction methods (Pearson r = 0.585, 0.618 and 0.621 for the network, CSD, and hybrid methods, respectively) (Supplementary Figure 5b, Bottom). These findings indicate that, although the tractography outputs differ qualitatively, they achieve broadly comparable performance in predicting tracer-defined structural connectivity. Notably, validation against tracers is currently limited to macaque datasets where tracer measurements are available and cannot readily be replicated in the human datasets.

### Superficial white matter and cortical-subcortical connectivity of the subthalamic nucleus

The observed differences in FOD structure motivated targeted tractography analyses in regions where the network’s properties may improve fibre reconstruction. Figure 6 highlights two example applications in which the network-derived FODs provide benefit: superficial white matter (SWM) reconstruction from single-shell dMRI data, and tracking through the subcortex, demonstrated here by mapping cortical connectivity to the subthalamic nucleus (STN).

**Figure 6.**
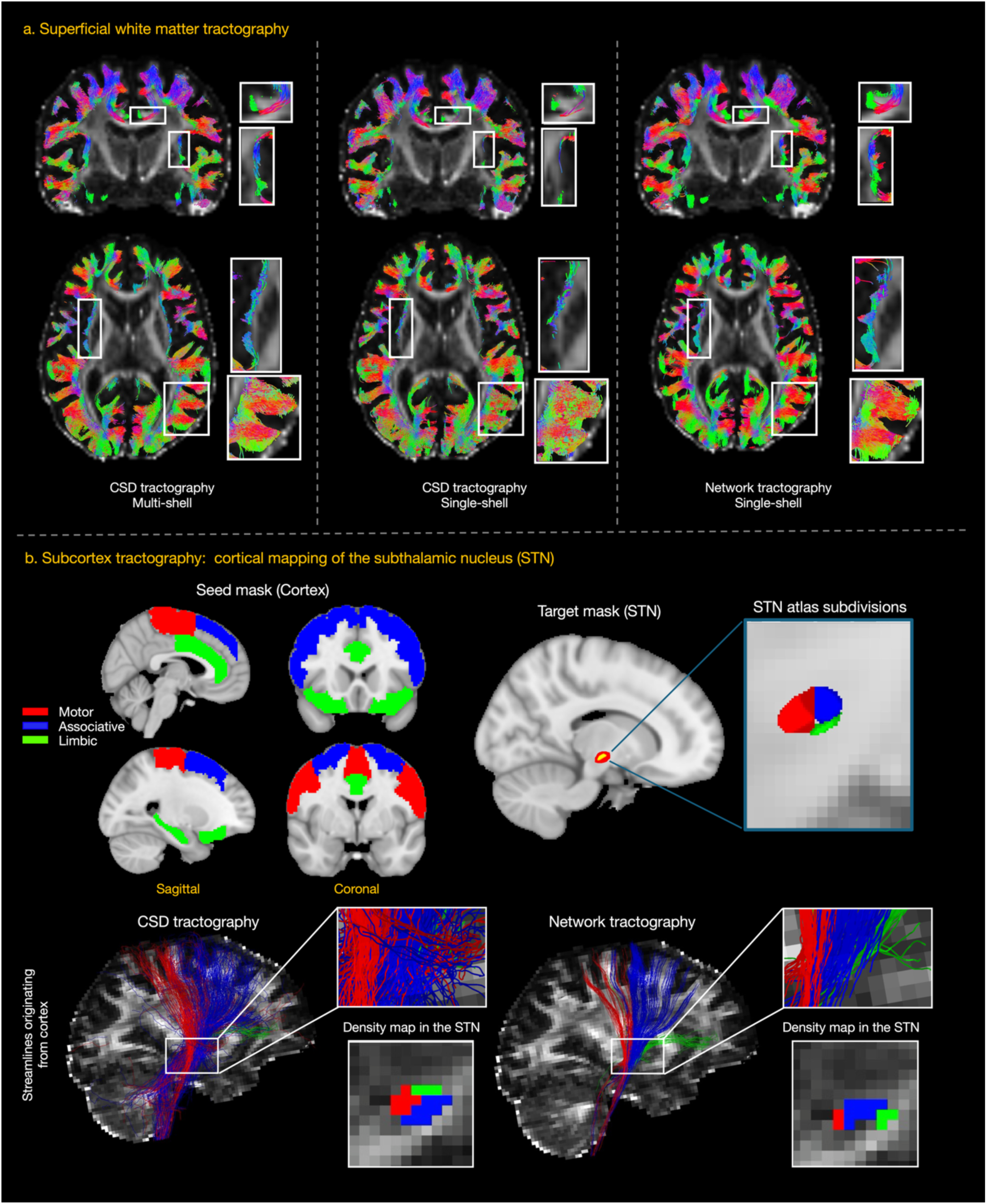
Tractography applications in SWM and subcortical tractography. a) Reconstruction of superficial white matter (SWM) in the human brain in the coronal (top) and axial (bottom) views. Multi-shell CSD tractography (left) serves as a reference for comparing single-shell CSD (middle) and network-derived single-shell tractography (right). In regions of high fibre curvature near the grey–white matter interface, single-shell CSD produces “noisy” or disjointed streamlines, whilst the network maintains better coherence and structural continuity. b) Topographical mapping of cortical to subthalamic nucleus (STN) connectivity. Cortical projections from motor (red), associative (blue), and limbic (green) regions are mapped to the STN. Network-derived tractography resolves a topographical organisation that aligns with established STN domains from the atlas^36^, whilst CSD tractography incorrectly places limbic projections in the superior domain.

Building on the network’s ability to resolve fibre orientations near grey matter, we evaluated its performance on superficial white matter (SWM), which consists primarily of short-range association fibres. SWM plays a critical role in local cortical communication, and its disruption is increasingly linked to various neurodevelopmental and neurodegenerative disorders^37^. SWM characterisation remains a significant challenge for dMRI tractography due to high fibre curvature, small fascicles, and proximity to the cortex (Figure 6a). Multi-shell CSD tractography from UK Biobank data was used as a reference and compared against tractography derived from (i) single-shell CSD and (ii) the network using the same single-shell input. In sulcal regions where single-shell CSD streamlines break or terminate prematurely due to the high curvature of U-fibres, the network robustly maintains continuity along the sulcus, following the trajectories observed in the multi-shell SWM. Moreover, in the posterior axial view in Figure 6a, the network FODs produce more coherent and organised pathways, whereas single-shell CSD yields disorganised and noisy streamlines.

Figure 6b examines cortical connectivity to the subthalamic nucleus (STN), a key clinical target for deep brain stimulation (DBS) in movement disorders such as Parkinson’s disease^38^. Accurate characterisation of cortical-STN connectivity is important because functional outcomes of stimulation depend critically on the specific cortical regions it manipulates, requiring sub-mm accuracy of DBS placement for ideal outcomes. For the STN, this includes motor, associative, and limbic circuits. However, reconstructing these pathways to the STN using dMRI is challenging due to the small size of the STN, its location deep within the brain, partial volume effects where tracking near and through subcortical grey matter is difficult due to the low anisotropy, and the presence of complex fibre configurations within surrounding subcortical white matter.

Given the increased orientational coherence observed in network-derived FODs within subcortical regions (Supplementary Figures 3 and 4), we investigated whether this translated into improved mapping of cortical projections (motor, associative and limbic) to the STN. Our results in Figure 6b demonstrate that the anatomical organisation derived from the network FODs aligns well, though imperfectly, with the established atlas for the STN sub-domains at 0.22 mm^36^. Specifically, the limbic projection is localised to the anteroinferior domain of the STN, the motor projection is more posterior, and the associative projection is situated in the anterosuperior domain. This topographical organisation is correctly resolved by the network-derived FODs. In contrast, the streamlines derived from CSD FODs are anatomically inconsistent, incorrectly placing limbic projections to the superior domain of the STN. Taken together, these results suggest the network may enable more precise DBS placement and improved patient outcomes.

## Discussion

In this study, we present a domain adaptation network to transfer microscopy-based information to in vivo dMRI, enhancing fibre orientation estimation in subjects where microscopy is not available. Our method is designed to process single-shell dMRI data, represented using spherical harmonics, making it applicable to a wide number of dMRI acquisitions without strict or heavy data requirements (only a single b-value and arbitrary gradient directions), facilitating broad application. Our results highlight the utility of cross-modal, cross-tissue and cross-species domain adaptation to transfer microscopy insights from animals to in vivo MRI in humans. Our microscopy-informed fibre orientations enable biologically meaningful fibre tracking, capturing internal white matter organisation, demonstrated by the internal capsule topography, and improving superficial white matter and cortical-subcortical tractography, including connectivity-based segmentation of the STN in vivo. The code and pre-trained network parameters are made publicly available for general application to single-shell datasets and future model development.

The network effectively maps dMRI signals to fibre orientations while preserving microscopy features. For example, the network-derived FODs successfully resolve patterns of internal white matter organisation within the internal capsule that mirror the topographical layout of the motor cortex, even in in vivo human data. This organisation has previously been demonstrated using invasive methods in the macaque^31,32^ and, in our prior work^27^, shown to arise from microscopy-informed hybrid FODs, on which the network is trained. Here, the network successfully transfers the FOD characteristics required to reproduce this pattern across both tissue state (postmortem to in vivo) and species (macaque to human). Our network FODs exhibit lower dispersion (local fanning) and fewer crossing fibres than CSD FODs, echoing observations from the PLI microscopy and hybrid FODs^23^. While the true number of crossing-fibre populations remains difficult to validate, given that no single microscopy modality provides an unambiguous ground truth for fibre organisation, our network consistently reproduces the microstructural characteristics of its PLI-derived training targets, highlighting both the specificity of the learned representations and the importance of microscopy choice in dataset design. These differences are particularly notable in the cortex, where the network FODs more closely capture gyral fanning of myelinated fibres, and cleaner branching patterns near the white-grey matter boundary, in line with myelin-sensitive PLI measurements on which the hybrid orientations are built.

This improved anatomical specificity resulted in superior single-shell superficial white matter (SWM) tractography. Reconstructing superficial fibres using diffusion tractography is challenging due to their small size, large curvature and proximity to grey matter and partial volume effects. These challenges are further exacerbated in single-shell dMRI, where the limited diffusion-weighting information constrains the ability to distinguish multiple tissue compartments and robustly resolve fibres in the cortex. By directly inferring microscopy-informed fibre architecture from the dMRI signal, without relying on fixed modelling assumptions or anatomical priors, the network mitigates some of the limitations of single-shell acquisitions. This highlights the network’s potential to bridge the gap between research-grade and clinically feasible dMRI protocols for SWM mapping.

It is instructive to consider what the network is learning. At a conceptual level, the network learns the relationship between fibre orientations and the dMRI signal, an implicit generalisation of the fibre response function (FRF). In conventional analyses, the diffusion signal is modelled as the convolution of the FOD with a response function. However, these quantities cannot be robustly estimated simultaneously voxel-wise, without considerable modelling constraints, from typical data. As a result, methods such as CSD first estimate a response function, from voxels assumed to contain a single fibre population, and then apply it as a fixed deconvolution kernel across the brain^14^. In single-shell data, only a single global response function is typically used, and even in multi-shell settings, the model remains constrained to a small number of predefined tissue types. However, microscopy-informed studies have shown that response functions likely vary substantially across the brain, and that enforcing a global response function can lead to systematic biases, including overestimation of fibre dispersion^30,39^. Rather than explicitly modelling this relationship, our approach learns it directly from data. The network implicitly captures spatially varying, tissue-specific mappings between diffusion signals and fibre orientations, without imposing a fixed parametric form. This flexibility provides a plausible explanation for its improved preservation of microscopy-derived features, particularly in regions of complex architecture such as superficial white matter. More broadly, it suggests that relaxing the assumption of a global response function may be key to improving fibre orientation estimation from clinically feasible dMRI.

Deep learning methods have previously been proposed to estimate fibre orientations from dMRI, similarly aiming to overcome the limitations of conventional biophysical models. Davood et al. demonstrated that a multilayer perceptron can effectively map from dMRI signals, interpolated onto a fixed spherical grid in q space, to FODs^40^; while Lucena et al. used a 3D convolutional neural network to predict “multi-shell” FODs from single-shell data, improving acquisition efficiency^41^. Microscopy-informed approaches have also been explored, with Zifei et al. training a network using tract-tracing data from the Allen Mouse Brain Connectivity Atlas to predict axonal connectivity^12^. However, the applicability of such approaches to the human brain remains limited by substantial differences in white matter organisation between the mouse and human^42^. In contrast, our approach integrates in vivo and postmortem dMRI with whole-brain microscopy within a single macaque brain, enabling direct learning of microscopy-informed fibre architectures in a model with human-like structural complexity. By combining matched in vivo and postmortem data with a domain adaptation framework, the network is able to transfer microstructural features across both tissue state (postmortem to in vivo) and species (macaque to human). Together, these advances provide a generalisable tool for integrating microscopy features into human connectivity research.

The study has several limitations. First, training was performed on data from a single macaque brain. Although this dataset contains a large number of voxels (>218,000 voxels for the postmortem dMRI) across multiple acquisitions, reliance on a single subject may increase the risk of overfitting. Expanding the training set to include additional brains with matched in vivo, postmortem, and microscopy data will likely improve robustness, as datasets of this nature become increasingly available.

Second, the training data were derived from a largely “healthy” brain (i.e., the animal did not die from neurological conditions), and evaluation was performed on non-clinical human datasets (HCP and UK Biobank). As such, the behaviour of the model in the presence of pathology, where diffusion signals may fall outside the training distribution, remains unknown. More broadly, the approach assumes that the relationship between fibre orientations and the diffusion signal learned from a healthy individual generalises across subjects, without explicitly modelling pathological variability.

Third, there was a resolution mismatch between the source and target domains during domain adaptation: the postmortem dMRI was acquired at 0.6 mm isotropic resolution, whereas the in vivo dMRI data were acquired at 1 mm. While the adaptation framework implicitly accounts for this discrepancy, the network architecture does not explicitly address the resolution differences, which may limit performance.

Finally, although our results are promising, validation in humans remains indirect: because no ground truth is available in vivo, we rely on anatomical plausibility rather than direct measurement. This highlights one of the key challenges in FOD reconstruction: how to know which outputs are “better”. Existing approaches compare dMRI FODs to microscopy “ground truths”, either through MRI-microscopy FOD comparisons at the voxel level or through downstream tractography with comparisons via tracers. Here our network, trained on microscopy, produces FODs more similar to the PLI, and comparable similarity to tracer-based connectivity via tractography. However, while the network shows clear benefits in some regions, microscopy-informed FODs should be interpreted with caution and require further validation. For example, the reduced prevalence of crossing fibres is consistent with PLI microscopy, but whether this reflects true biology remains uncertain. Future work could assess clinical relevance through post hoc analyses as a form of validation, for example by evaluating whether improved connectivity estimates better predict outcomes following interventions such as deep brain stimulation.

Several directions remain for future work. We adapted our network to dMRI at 1.25 mm resolution from HCP data and 2 mm resolution from UK Biobank data, both with b = 1000 s/mm², subsampled from multi-shell acquisitions. Future work should extend our approach to varying diffusion acquisition protocols such as sparse acquisition protocols suitable for pre-operative tractography^43^, and validate performance across larger populations to enhance robustness and broader applicability.

In summary, we developed a domain adaptation network to translate fibre information learnt from microscopy into in vivo dMRI. This data-driven approach estimates fibre orientations without relying on complex biophysical modelling, recovering meaningful microscopy-informed features and demonstrating clear benefits for precision fibre tracking. The potential improvements in in vivo tractography, including in clinically relevant pathways, highlight promising future applications in interventions such as connectivity-informed targeting for brain stimulation. By enabling the transfer of microstructural information across tissue state and species, this work suggests a general paradigm for integrating invasive microscopy with non-invasive human imaging.

## Methods

### Data: macaque MRI and microscopy

This study used data from two macaque datasets: the BigMac dataset, combining in vivo and postmortem MRI with whole-brain microscopy for training (with held-out validation and test data), and an independent in vivo macaque MRI dataset used exclusively for testing.

### The BigMac dataset

The BigMac dataset was acquired and pre-processed as described in Howard et al. 2023^8^. An adult rhesus macaque brain was scanned both in vivo and postmortem and then processed for whole-brain microscopy.

In vivo MRI was acquired on a 3T whole-body scanner, including 0.5 mm isotropic structural images using a MP-RAGE sequence and diffusion-weighted images using spin-echo echo-planar imaging (SE-EPI): echo time (TE) = 100 ms, repetition time (TR) = 8.2 s, resolution = 1 mm isotropic, b-value = 1000 s/mm², 1100 diffusion gradient directions, including 81 unique gradient directions and 144 volumes with negligible diffusion weighting (b∼0 s/mm²).

Postmortem MRI was acquired on a 7T preclinical scanner, including 0.3 mm isotropic structural images using a multi-gradient echo sequence (MGE 3D) and diffusion-weighted images with two b values: 1) b=4000 s/mm² at 0.6 mm isotropic resolution with 128 gradient directions and 8 volumes with negligible diffusion weighting: TE = 25.4 ms, TR=10 s; δ/Δ = 7/13 ms and 2) b=10,000 s/mm^2^ at 1.0 mm isotropic resolution with 1000 gradient directions and 40 b∼0 s/mm² volumes: TE = 42.4 ms, TR= 3.5 s; δ/Δ = 14/24 ms.

The structural images were used for co-registration and mask generation for tractography, whilst the dMRI was used for fibre orientation construction via our network and CSD. All data were pre-processed, including susceptibility, eddy current and motion correction as appropriate^8^.

After scanning, the postmortem brain was sectioned along the anterior-posterior axis with consecutive slices assigned to one of six microscopy contrasts. This work uses the polarised light imaging (PLI) data from sections with a thickness of 50 µm repeated every 350 µm. PLI utilises the birefringence of myelin to estimate the primary fibre orientation per pixel^44,45^, here acquired at a resolution of 4 µm/pix. Precise co-registration of MRI-Microscopy was performed using purpose-designed software TIRL (v3.6)^46^.

### Hybrid dMRI-microscopy orientation generation

Hybrid dMRI-microscopy FODs were reconstructed in the BigMac dataset by combining information from dMRI and microscopy, following our previously described method^27^. PLI provided in-plane fibre orientations, while postmortem dMRI, modelled with the ball and stick approach (BAS)^13^, was used to approximate through-plane angles. Integrating these sources yielded high-resolution 3D fibre orientations at the spatial scale of the microscopy data.

These orientations were then aggregated across local neighbourhoods corresponding to individual MRI voxels to generate hybrid FODs, which served as the “ground truth” target during network training. To match the MRI data used for training, hybrid FODs were reconstructed at both 0.6 mm and 1 mm isotropic resolution in the postmortem dMRI space. The 1mm hybrid FODs were then registered to the BigMac in vivo dMRI space using the MRtrix3 command mrregister, and the FODs were reoriented according to the corresponding warp field^47^.

### Other macaque MRI

In vivo dMRI from another macaque brain was previously acquired using the same protocol as the BigMac in vivo dMRI session. This preprocessed data is openly available as part of the Oxford PRIME-DE dataset^48^. This data was solely used for network testing.

### Data: in vivo human MRI

Our method was evaluated on two openly available human MRI datasets: the UK Biobank, with 2 mm isotropic dMRI, and the Human Connectome Project (HCP), with 1.25 mm isotropic voxels. The UK Biobank protocol reflects data typically achievable in large-scale clinical studies (∼7 min scan time), whereas the HCP provides high-quality, research-grade data acquired with a longer scan duration (∼1hr).

### HCP

MRI for a single subject (female, <30 years) was obtained from the young adult cohort of the WU-Minn Human Connectome Project (HCP)^28^. dMRI was acquired on a 3T Siemens scanner using SE-EPI: TR = 5520 ms, TE = 89.5 ms, resolution = 1.25 mm isotropic, *δ*/Δ = 21.4/45.5 ms, b-values of 1000, 2000 and 3000 s/mm² with 30 unique directions per shell repeated with reversed phase-encoding polarity (180 in total) and 36 interspersed b∼0 s/mm² volumes. Only the single-shell b = 1000 s/mm² volumes and b∼0 s/mm² volumes were used for input to the network. The data were previously pre-processed as described^11^ including susceptibility, eddy current and motion correction.

T1-weighted structural MRI - used for co-registration only - was acquired at 0.7 mm isotropic resolution using a MPRAGE sequence. Registrations were performed using FSL FLIRT/FNIRT^49,50^.

### UK Biobank

MRI data for two subjects were accessed from the UK Biobank^29^. dMRI was acquired on a 3T Siemens scanner using a SE-EPI: TR = 3600 ms, TE = 92 ms, resolution = 2 mm isotropic, *δ*/Δ = 21.4/45.5 ms, b-values of 1000 and 2000 s/mm² with 50 directions per shell, 5 b∼0 s/mm² volumes and 3 blip-reversed b0 volumes for distortion correction, resulting in a ∼7 minutes scan time. The data were previously pre-processed as described^29^ including susceptibility, eddy current, and motion correction. Only the single-shell b = 1000 s/mm² and b∼0 s/mm² volumes were used for input to the network. Diffusion data were processed using constrained spherical deconvolution (CSD), as described below.

T1-weighted structural MRI was acquired at 1 mm isotropic resolution using a 3D MPRAGE sequence. T1-weighted scans were processed using FreeSurfer^51^ to reconstruct pial and white matter surfaces. Volumetric registrations were performed using FSL FLIRT/FNIRT^49,50^.

### Constrained spherical deconvolution

Fibre orientation distributions were computed from single-shell dMRI in all datasets implemented in MRtrix3 (v3.0.3)^47^. The white matter response function was derived using the tournier algorithm, and FODs were subsequently reconstructed using constrained spherical deconvolution (CSD)^14^. The resulting FODs were compared with those predicted by the network.

We additionally processed the full multi-shell Biobank dataset using the dhollander algorithm to estimate tissue-specific response functions for white matter, grey matter, and cerebrospinal fluid (CSF)^15^. These response functions were then used within the multi-shell, multi-tissue (MSMT) CSD model to compute the FODs which were used for superficial white matter tractography.

### Datasets for network training, validation and testing

The network was first trained using macaque in vivo MRI (1 mm, b = 1000 s/mm²) and postmortem MRI (0.6 mm, b = 4000 s/mm²) dMRI as inputs, with co-registered hybrid FODs as targets at the same resolution. The training dataset was partitioned into training (80%) and validation (20%) sets, with voxels outside the brain mask excluded.

Unseen data were then used to evaluate the network’s generalisability and performance. This included: (1) postmortem dMRI from the same macaque at 1.0 mm isotropic resolution, b=4000 s/mm²; and (2) in vivo dMRI from another macaque.

The network was subsequently adapted for in vivo human MRI using two datasets: single-shell HCP and Biobank dMRI data, both b=1000 s/mm². The unsupervised adaptation process for human data did not require FOD targets, as described below.

Spherical harmonics of order 8 were fitted to the normalised diffusion-weighted signal (S/S0) prior to input to the network, where S_0_ represents the non-diffusion-weighted signal with b∼0 s/mm². Brain volumes were then subdivided into 3×3×3 voxel patches using a sliding window approach (stride = 1), to provide local spatial context. The input data were standardised to zero mean and unit variance prior to model training.

### The network

#### Network architecture

We developed a domain adaptation network with three components: a feature extractor, a predictor and a discriminator^52^ (Figure 2). The feature extractor and predictor generated FODs from dMRI, while the domain discriminator ensured invariance between (i) postmortem and in vivo data, and (ii) human and macaque input data. The network received small blocks of MRI data (3×3×3 voxels) as input and produced the microscopy-informed FOD for the central voxel. By processing cubes of voxels through convolutional layers, rather than individual voxels, the network incorporated local neighbourhood information to constrain the output FOD since fibre orientations are generally spatially continuous. The predictor and domain discriminator were implemented using fully connected layers. Within the predictor, the initial fully connected layer was followed by Batch Normalisation, ReLU activation, and Dropout. Subsequent hidden layers utilised ReLU activation, whereas the final output layer utilised a linear activation to output spherical harmonic coefficients. During training, the predicted FOD was directly compared to the hybrid dMRI-microscopy FOD from the same voxel^27^, which was taken to be the ground truth target.

Both the input dMRI and output FOD were represented using spherical harmonics. By using this representation, the diffusion signal was described by a fixed number of spherical coefficients (45 in this study) rather than an arbitrary number of gradient directions. This allows the network to be applied to existing datasets without strict acquisition requirements regarding the number of gradient directions. Specifically, we require the input data to have at least 45 gradient directions to support the fitting of the 45 spherical harmonic coefficients, but the network could be readily retrained with a lower order spherical harmonic basis to accept data with fewer gradient directions. The output FOD was also represented using spherical harmonics, as is common in dMRI methods such as constrained spherical deconvolution. This approach provides the FOD in a standard format compatible with existing tractography algorithms and eliminates the need to predefine the number of fibre populations per voxel, as spherical harmonics can describe a free-form orientation distribution on the sphere.

The discriminator was designed as a binary classifier terminating with a softmax layer, providing a probability distribution over the source and target domains. The discriminator facilitated domain adaptation in two contexts. The network was first trained with postmortem (source) and in vivo MRI (target) data, using domain adaptation to enforce tissue-state invariance. It was then re-trained using macaque (source) and human MRI (target) data, employing the same strategy to ensure species invariance. For domain adaptation, we first trained the discriminator to determine whether the input data originated from the source or target domain and then used adversarial training to confuse the discriminator. This forced the feature extractor to learn a latent representation that was domain-agnostic (i.e., invariant to tissue state or species). This procedure ensured that the network trained on postmortem macaque data could later be applied to in vivo human data without requiring matched microscopy.

#### Network training: macaque data only

The network was trained via a two-stage adversarial approach inspired by Dinsdale et al. 2021^25^.

In the first stage, we used an alternating optimisation scheme to decouple the main task of fibre orientation estimation from domain discrimination. The feature extractor and predictor were optimised to minimise the error of fibre orientation estimation, while the domain discriminator was trained to distinguish between in vivo and postmortem MRI inputs. Crucially, when updating the feature extractor and predictor, the weights of the domain discriminator remained frozen, and vice versa, ensuring stable convergence of the adversarial components.

The main task loss function was defined as the mean squared error (MSE) between the amplitudes of the predicted and ground truth FODs. Each FOD was first converted from spherical harmonics to amplitudes across the sphere (along 256 directions) prior to MSE calculation. Spatial misalignment between MRI and microscopy data could result in slight rotational discrepancies between the predicted FOD relative to the ground truth. To address this, we implemented a rotation-compensated loss function: the predicted FODs were rotated about the x, y and z axes by a small range of angles (<10 degrees on each axis). Each rotated prediction was then compared to the hybrid FOD ground truth, and the final loss was defined as the minimum mean square error across all permitted rotations. This approach ensured that minor angular misalignment due to registration discrepancies between MRI and microscopy was accounted for during training. The loss was defined as follows, where ℛ denotes the set of permitted rotations, *R* a single rotation from this set, ŷ the predicted FOD, y the ground truth FOD and N the number of sampled directions (N=256).

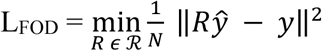

The domain discriminator was trained using cross-entropy loss with one-hot encoded domain labels ([0,1] for source and [1,0] for target). Minimising the cross-entropy loss maximised the discriminator’s ability to distinguish between the source and target domains.

In the second stage, the domain discriminator parameters were frozen, and adversarial training was applied by introducing a domain confusion loss, calculated as the Shannon entropy of the discriminator’s softmax output. By backpropagating the gradients from this loss to the feature extractor, the model was trained to produce domain-agnostic features, such that the discriminator’s softmax output approaches a uniform distribution across domain labels. This effectively removed domain-specific information. To preserve accurate fibre orientation estimation, the feature extractor and predictor were optimised using a joint objective comprising the confusion loss and MSE loss:

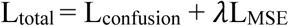

where *λ* is a hyperparameter that controls the trade-off between FOD estimation accuracy and domain invariance (*λ* = 0.05). L_confusion_ is the Shannon entropy, defined as -∑ *p_j_* log(*P_j_*) where *p_j_* represents the probability of the input belonging to domain j.

This two-stage adversarial framework thus balanced two opposing objectives: the domain loss encouraged the discriminator to distinguish between in vivo and postmortem MRI inputs, while the confusion loss encouraged the feature extractor to eliminate these differences, ensuring domain invariance in the learned representations. The hyperparameters were defined by a learning rate of 1e-4 and a batch size of 128, using the Adam optimiser; training lasted 10 epochs during stage 1 and 10 epochs during stage 2.

#### Network training: adaptation to human data

We adapted our macaque-trained network to human in vivo data using an unsupervised domain adaptation (UDA) framework. The goal was to establish species invariance between macaque and human dMRI inputs. The UDA employed a two-stage adversarial training strategy similar to that described above, but here adapted for macaque MRI (source domain) with paired hybrid FODs as ground truth (“labelled macaque data”), and human MRI (target domain) without corresponding ground truth FODs (“unlabelled human data”).

Labelled macaque data provided supervision for fibre orientation estimation, while unlabelled human data guided the network to adapt to interspecies differences by exposing the network to source-target variation. In the first stage, the feature extractor and predictor were trained on labelled macaque data to predict microscopy-informed FODs, while the discriminator was trained in alternation to distinguish between human and macaque dMRI inputs.

In the second stage, the discriminator parameters were frozen, and the feature extractor and predictor were trained using a combined loss that balanced two objectives: 1) promoting species invariance through domain confusion loss, and 2) maintaining accurate FOD estimation via the MSE loss.

#### Network FODs post-processing

Network-derived FODs were normalised on a voxel-wise basis using their zeroth-order spherical harmonic (*l*=0) coefficient scaled by 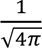, such that the integral of the FOD over the unit sphere was normalised and thus comparable across methods, facilitating visual comparison and tractography analysis.

### Macaque Tractography

#### White matter organisation in the internal capsule

We examined white matter organisation within the internal capsule (IC) building on our previous work using hybrid fibre orientations^27^. Streamlines seeded from functionally defined motor cortex subregions (trunk, arm, and face) were used to assess whether network-derived fibre orientations preserved the known anterior–posterior topographical organisation of projections through the internal capsule^31,32^, as also previously demonstrated using hybrid FODs but not CSD tractography. Motor cortex ROIs were manually drawn on the BigMac structural MRI surface, converted to volume space, and warped to the in vivo and postmortem dMRI spaces for use as tractography seed masks. The internal capsule mask was obtained from the D99 atlas v2.0^21^ and similarly warped to both postmortem and in vivo BigMac diffusion spaces.

Tractography was performed using MRtrix3 and the iFOD2 algorithm^14^, seeding from the ROIs in the cortex and only retaining streamlines through the internal capsule target mask. Tracking parameters included 30 seeds per voxel, with a step size of 0.1 mm, a cut-off value of 0.05, and streamline length constraints of 2-80 mm. Tract density maps were generated for each cortical ROI, with densities normalised by the total number of streamlines originating from each ROI, and voxels within the internal capsule were colour-coded according to which ROI contributed the highest streamline density. Outputs were compared between running tractography with the network-, hybrid-, and CSD-based FODs.

#### U-shape fibre tractography

U-shape fibres are short association fibres that connect neighbouring gyri with high curvature. In macaques, these fibres were tracked using cortical surface masks from neighbouring gyri which were manually delineated at the grey matter-white matter boundary within dMRI space. Tractography, using the parameters described above, was seeded from one gyrus and only streamlines that tracked to the target mask in the neighbouring gyrus were retained.

#### Comparison to retrograde tracer connectivity

We compared connectivity matrices derived from our network- and CSD-based tractography with a weighted connectivity matrix derived from retrograde tracer data, used here as a ground truth estimate of structural connectivity^33–35^. The tracer-derived matrix, openly available on the Brain Analysis Library of Spatial maps and Atlases (BALSA) database^53^, was constructed from tracer injections into 29 regions of 91 cortical areas of the macaque left hemisphere. Connectivity strength was quantified as the fraction of labelled neurons (FLNe), defined as the proportion of labelled neurons in each target area relative to the total number of labelled neurons at the injection site. To exclude very sparse tracer connections that are unlikely to be captured by tractography, connections with FLNe < 0.0001 in the tracer data were removed. The macaque cortical atlas was co-registered to BigMac postmortem dMRI, using warp fields generated by FSL’s FLIRT/FNIRT. Tractography was then seeded from the 29 cortical injection sites using the parameters described above, and the number of streamlines projecting to each of the 91 cortical regions, normalised by the total number of streamlines originating from each injection site, was used to construct the 29 × 91 weighted connectivity matrices for both network- and CSD-derived FODs.

The connectivity matrix was evaluated via receiver operating characteristic (ROC) analysis, using the tracer connectivity matrix as the gold standard. The area under the curve (AUC) was calculated to determine the specificity and sensitivity of each reconstruction method (connectivity derived from network, hybrid, and CSD FODs in the postmortem macaque). We further quantified the false positive rate (FPR) at a tracer threshold of 0.0001 to evaluate the anatomical validity of the identified connections. Finally, we performed correlation analyses between the tracer connectivity matrix and the normalised connectivity matrices from both the network and CSD FODs. Log-log correlation plots were generated, and Spearman’s correlation (*ρ*) used to quantify the agreement between the tractography-derived estimates and the tracer ground truth.

### Human tractography

#### Major white matter tract reconstruction

Whole-brain probabilistic tractography was performed using FODs generated by both the network and CSD with the iFOD2 algorithm. Tracking parameters included 30 seeds per voxel, a step size of 0.2 mm, a cutoff value of 0.1, and streamline length constraints of 5-120 mm. Tracts were segmented using inclusion and exclusion masks from the XTRACT atlas^54^.

#### White matter organisation in the internal capsule

The topographical organisation of the human internal capsule was evaluated using UK Biobank data, following the same tractography protocols described for the macaque. Cortical seed masks for the trunk (A4t, area 4), arm (A4ul, area 4) and face (A4hf, area 4) were defined in the MNI space derived from the Human Brainnetome Atlas, a parcellation atlas based on both anatomical and functional connections^55^. The anatomical mask for the internal capsule was obtained from the JHU ICBM-152 atlas^56^. Streamlines projecting from the cortical regions were tracked to the internal capsule target mask, and voxel-wise winner-take-all maps were generated based on the cortical ROIs with the highest streamline density.

#### Crossing fibre ratio

To evaluate the number of crossing fibres, we converted the FODs to discrete peaks using sh2peaks from MRtrix3^47^, identifying up to three principal fibre orientations per voxel with a threshold of 0.1, and calculated the number of distinct peaks per voxel. The prevalence of discrete fibre populations was quantified for hybrid and CSD FODs in the postmortem macaque, and for network FODs and CSD FODs in the UK Biobank human data.

#### Superficial white matter tractography

The superficial white matter (SWM) was reconstructed from UK Biobank data using a validated pipeline comprising two sequential steps: whole-brain tractography and SWM-specific filtering^57^.

Whole-brain probabilistic tractography was first performed by seeding from the grey-white matter interface generating 2 million streamlines with streamline propagation constrained by a white matter mask and parameters adapted for short fibres characteristic of SWM. The maximum streamline length was limited to 40 mm to exclude long-range fibres.

The tractogram was then iteratively filtered to isolate SWM fibres. First, a grey-grey matter filter retained only streamlines originating and terminating within cortical grey matter.

Second, a hemisphere-hemisphere filter removed inter-hemispheric commissural fibres by restricting streamlines to a single hemisphere. Third, a white matter filter ensured streamlines traversed white matter, excluding superficial U-fibres confined to grey matter. This multi-stage filtering approach, validated in prior work^57^, enhances specificity to SWM while improving cortical coverage.

Cortical surfaces and tissue boundaries, derived from T1-weighted structural images using FreeSurfer, served as anatomical masks to remove any streamlines that did not meet the above criteria^51^. The pipeline was applied to FODs generated by both single- and multi-shell CSD and the network to enable comparative analysis of SWM reconstructions.

#### Cortical -STN Tractography

The topographical projections from the cortex to the subthalamic nucleus (STN) were reconstructed using the UK Biobank dataset. Cortical seed masks representing associative, motor, and limbic regions were derived from the Harvard-Oxford cortical atlases (Makris 2006 parcellation)^58^. The STN and its corresponding subregions were defined using the DBS Intrinsic Template Atlas (DISTAL) in MNI space and the STN mask was utilised as an inclusion mask for streamline selection^36^. All anatomical masks were registered to the individual dMRI space.

Following the probabilistic tractography protocol described above, streamlines were seeded from the cortical ROIs and constrained to those intersecting the STN target mask. Voxel-wise tract density maps were generated for each cortical ROI and normalised by the total streamline count. To visualise the topographical organisation within the STN, voxels were colour-coded using a winner-take-all approach, assigned to the cortical ROI contributing the highest normalised streamline density. This mapping was performed for both CSD and network-derived FODs.

## Acknowledgement

SZ is supported by the Chinese Government Scholarship and NIH (Grant UM1NS132207) and the Center for Mesoscale Connectomics. KLM, SJ, and AFDH were supported by the Wellcome Trust (224573/Z/21/Z, 215573/Z/19/Z and 221933/Z/20/Z). The Centre for Integrative Neuroimaging was supported by core funding from the Wellcome Trust (203139/Z/16/Z and 203139/A/16/Z). For the purpose of open access, the author has applied a CC BY public copyright licence to any Author Accepted Manuscript version arising from this submission.

