## Supplementary Figures for "Microscopy-informed structural connectivity mapping in the in vivo human brain via domain adaptation"

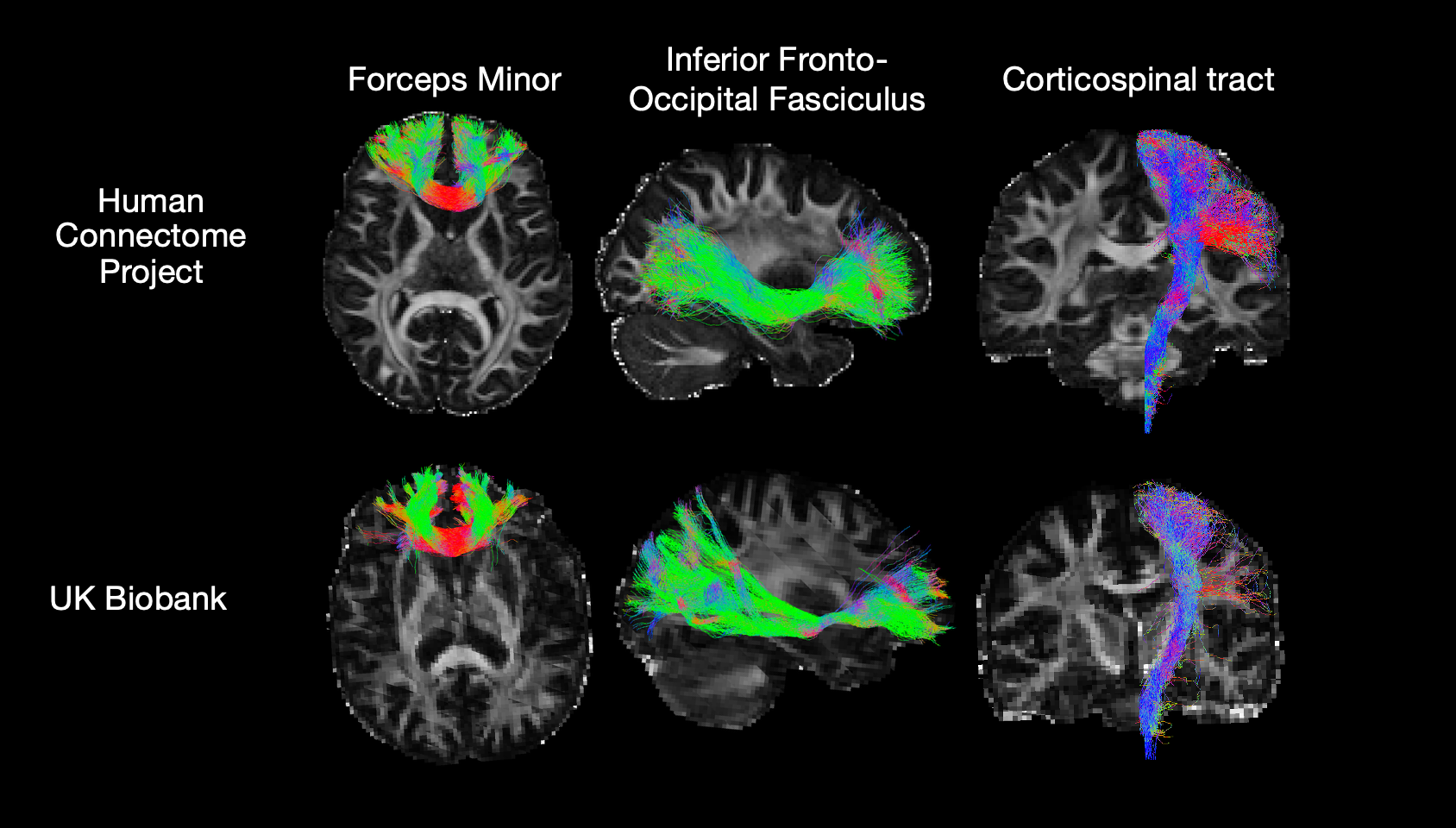

**Supplementary Figure 1: Major white matter tracts from CSD FODs.**

Example reconstructions of major white matter pathways derived from CSD FODs in datasets from the Human Connectome Project (top) and the UK Biobank (bottom). Three representative tracts are illustrated: the forceps minor, the inferior fronto-occipital fasciculus (IFOF), and the corticospinal tract (CST). These examples provide a direct comparison to the network-estimated reconstructions shown in Figure 4a.

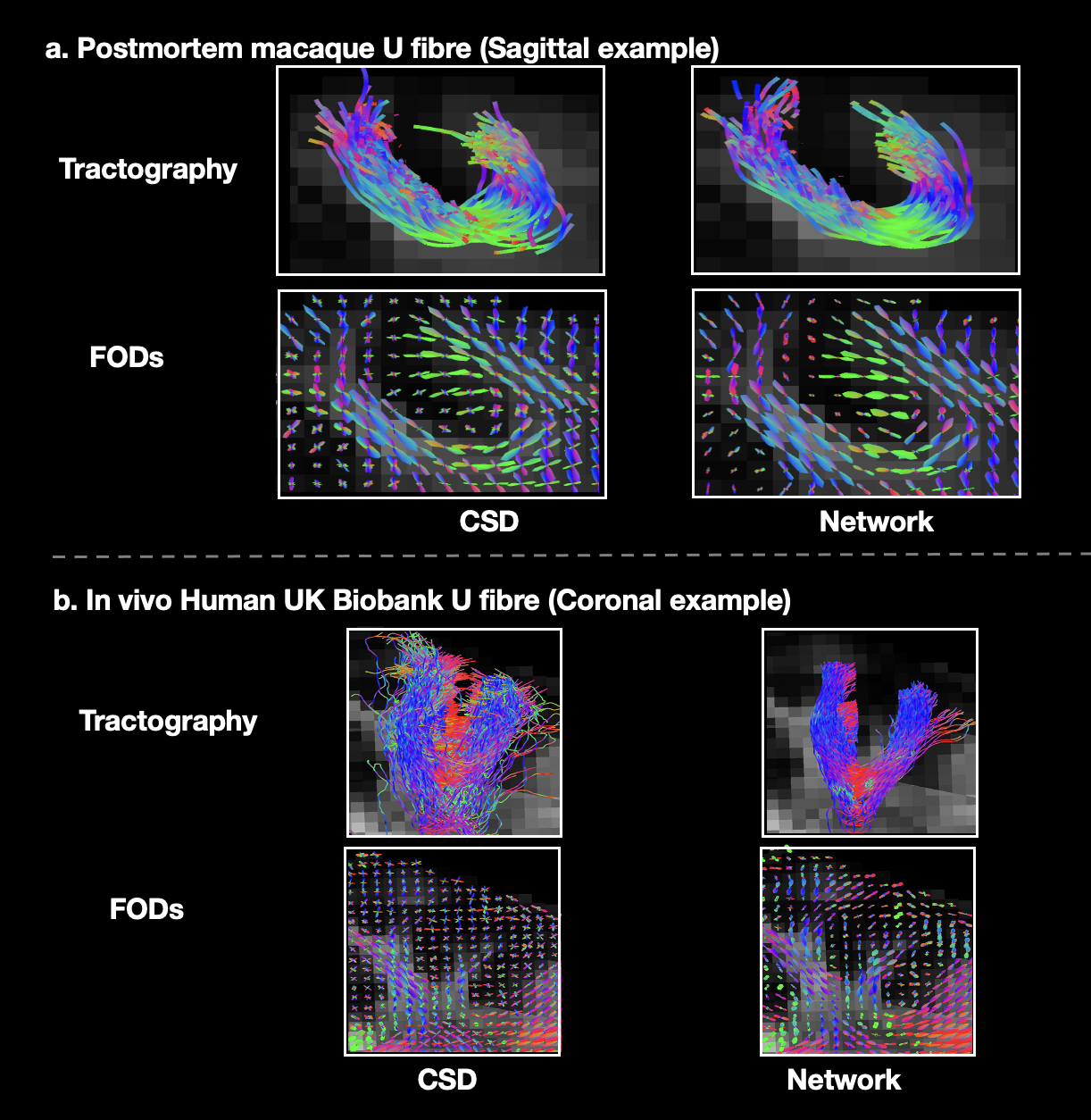

**Supplementary Figure 2: U-fibres tractography and FODs.**

U-fibre reconstructions using network and CSD FODs in postmortem macaque and in vivo human data. a) A macaque region where both CSD and network produce comparable U-fibre reconstructions. b) A UK Biobank example where CSD FODs yield noisy U-fibres, whereas the network delineates a coherent U-fibre. The difference reflects improved spatial continuity in network-estimated FODs near the cortical boundary, compared with noisier, less coherent CSD FODs across adjacent voxels.

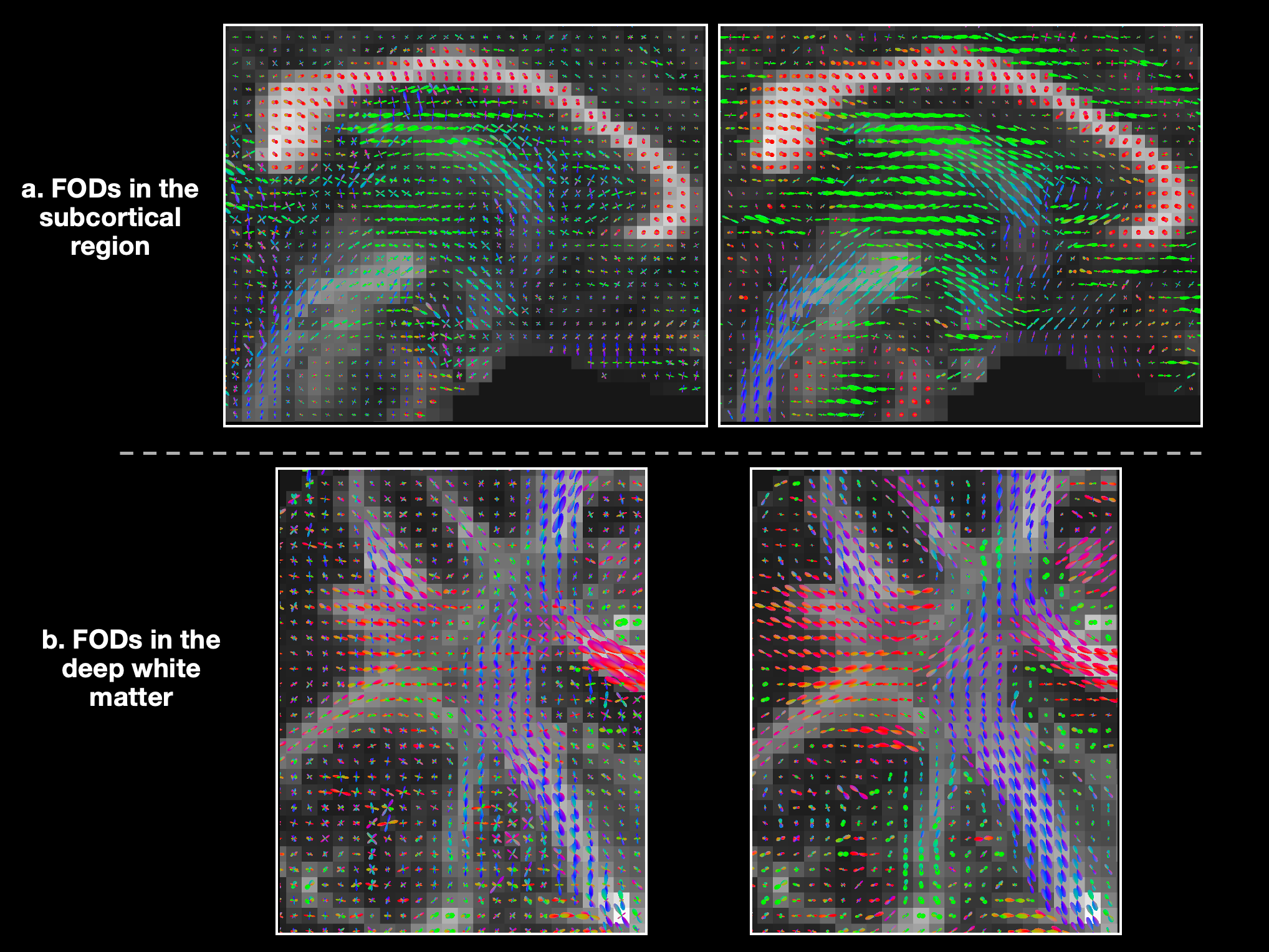

**Supplementary Figure 3: Comparison of CSD and network-estimated FODs across brain regions in UK Biobank data.**

FODs estimated using CSD (left column) and the network (right column) are shown in two anatomical regions: a) in the subcortical region, and b) in deep white matter.

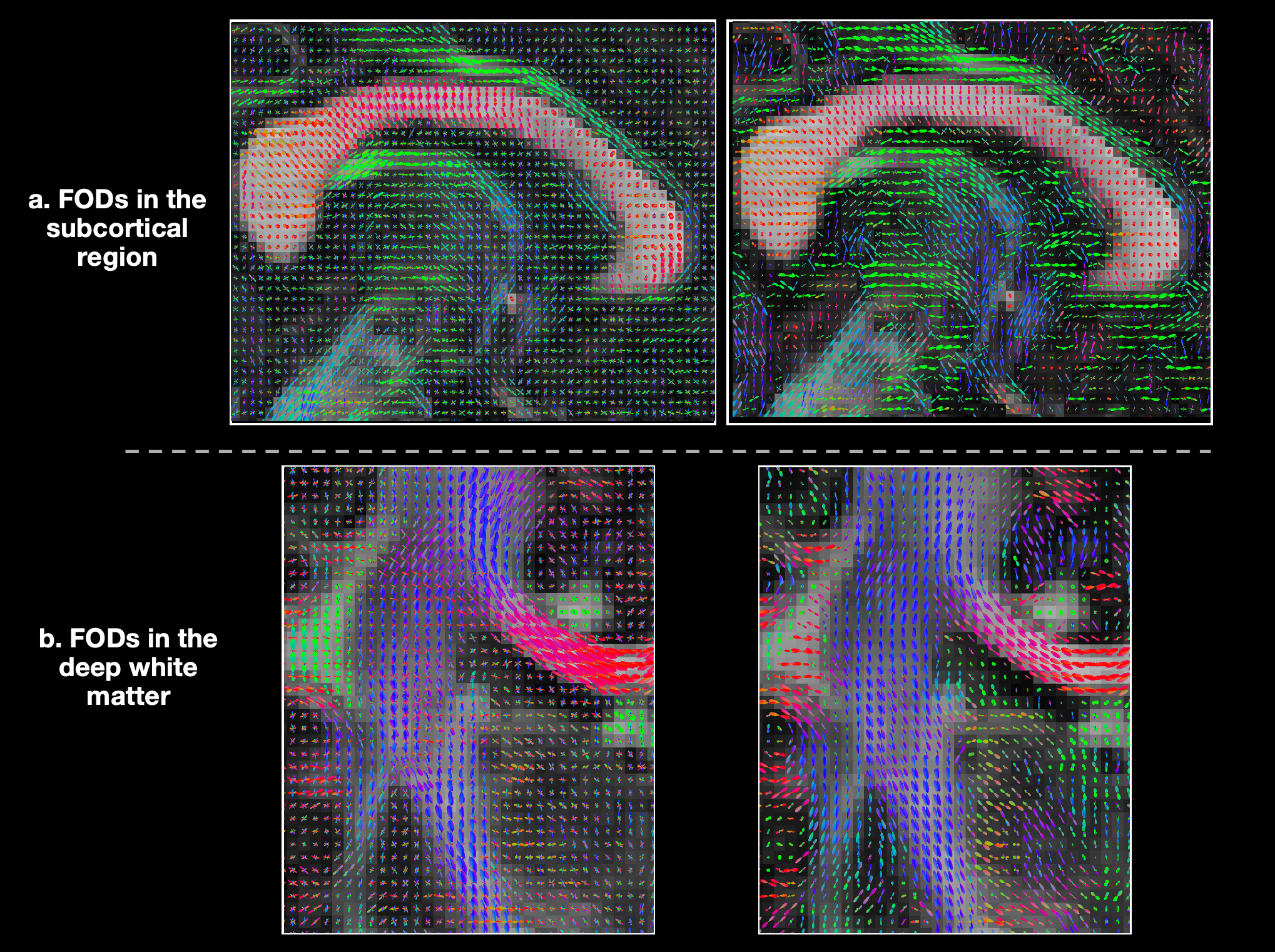

**Supplementary Figure 4: Comparison of CSD and network-estimated FODs across brain regions in HCP data.**

FODs estimated using CSD (left column) and the network (right column) are shown in two anatomical regions: a) in the subcortical region, and b) in deep white matter.

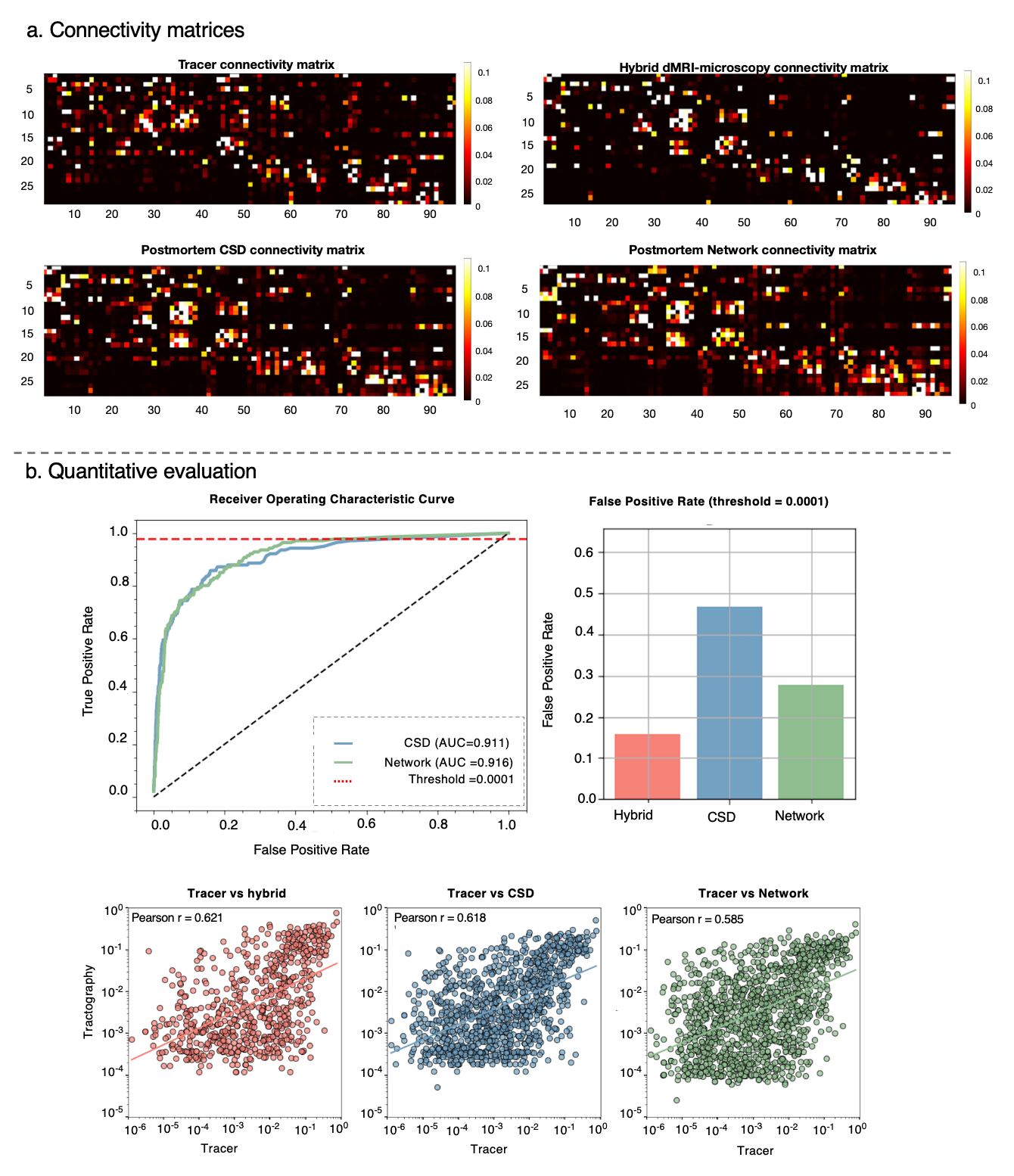

**Supplementary Figure 5: Tracer analysis.**

a) Connectivity matrices derived from tracer data, hybrid dMRI–microscopy FODs, postmortem CSD FODs and postmortem network FODs. b) Quantitative validation against retrograde tracer data. ROC analysis (top) and false positive quantification show slightly improved tract tracing accuracy (AUC = 0.916 for network and AUC=0.911 for CSD). Correlation plots (bottom) demonstrate that network-derived connectivity weights (r = 0.585) are comparable to those from CSD (r=0.618) and hybrid FODs (r=0.621) when compared to the tracer.
